# ALG-097111, a potent and selective SARS-CoV-2 3-chymotrypsin-like cysteine protease inhibitor exhibits *in vivo* efficacy in a Syrian Hamster model

**DOI:** 10.1101/2021.02.14.431129

**Authors:** Koen Vandyck, Rana Abdelnabi, Kusum Gupta, Dirk Jochmans, Andreas Jekle, Jerome Deval, Dinah Misner, Dorothée Bardiot, Caroline S. Foo, Cheng Liu, Suping Ren, Leonid Beigelman, Lawrence M. Blatt, Sandro Boland, Laura Vangeel, Steven Dejonghe, Patrick Chaltin, Arnaud Marchand, Vladimir Serebryany, Antitsa Stoycheva, Sushmita Chanda, Julian A. Symons, Pierre Raboisson, Johan Neyts

## Abstract

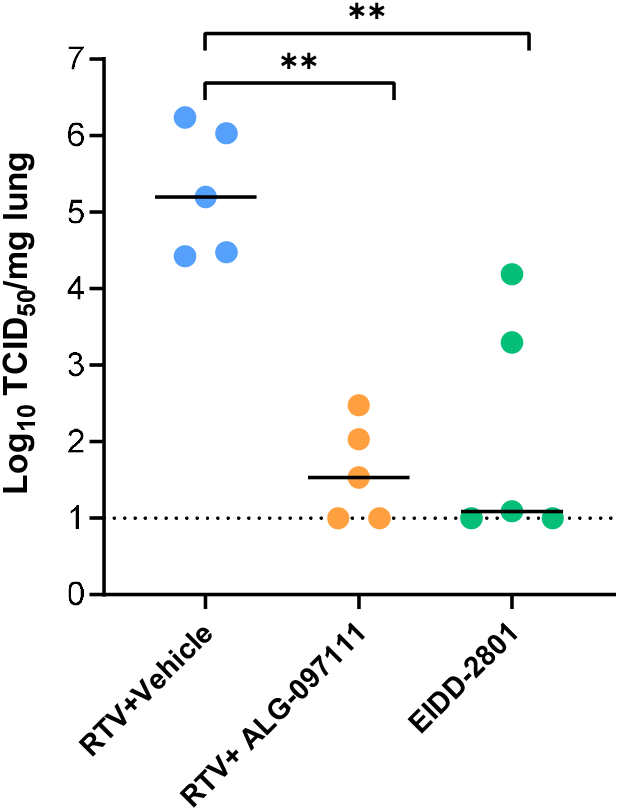

There is an urgent need for antivirals targeting the SARS-CoV-2 virus to fight the current COVID-19 pandemic. The SARS-CoV-2 main protease (3CLpro) represents a promising target for antiviral therapy. The lack of selectivity for some of the reported 3CLpro inhibitors, specifically versus cathepsin L, raises potential safety and efficacy concerns. ALG-097111 potently inhibited SARS-CoV-2 3CLpro (IC_50_ = 7 nM) without affecting the activity of human cathepsin L (IC_50_ > 10 μM). When ALG-097111 was dosed in hamsters challenged with SARS-CoV-2, a robust and significant 3.5 log_10_ (RNA copies/mg) reduction of the viral RNA copies and 3.7 log_10_ (TCID50/mg) reduction in the infectious virus titers in the lungs was observed. These results provide the first in vivo validation for the SARS-CoV-2 3CLpro as a promising therapeutic target for selective small molecule inhibitors.

## Introduction

Inhibition of the HCV and HIV viral proteases with antiviral drugs is a proven successful therapeutic approach. [1] With the continuing global COVID-19 pandemic and the urgent need for effective SARS-CoV-2 antivirals, investigators have repurposed previously discovered antiviral drugs initially designed against other viruses. These efforts have resulted in the approval of remdesivir, a nucleotide analogue inhibiting the nsp12 polymerase, previously developed for Ebola [2,3]. A second nsp12 inhibitor, MK-4482/EIDD-2801, is currently being evaluated in the clinic [4] [5]. The SARS-CoV-2 main protease (Mpro, 3CLpro) represents another promising target for antiviral therapy with no human homologue. Currently, the most advanced drug candidate targeting the SARS-CoV-2 3CLpro, is PF-07304814, a phosphate prodrug of PF-00835231 [6], discovered more than 15 years ago in the context of the SARS-CoV-1 outbreak [7]. Since the initial SARS-CoV-2 outbreak, several reports of 3CLpro inhibitors with *in vitro* potency against SARS-CoV-2 have been reported. [8] However, the lack of selectivity of some of these compounds, specifically versus cathepsin L, raises questions on the overestimation of the 3CLpro component of *in vitro* cellular potency and its translation towards *in vivo* activity. [9] Cathepsin L is involved in the entry of SARS-CoV-2 into host cells and therefore contributes to non-selective 3CLpro inhibitor potency in commonly used cell lines. GC-376, a broad spectrum inhibitor of 3C, 3C-like proteases and cathepsin L, failed to show a clear antiviral effect in monotherapy in a mouse model of SARS-CoV-2 infection.[10] In another study modest antiviral activity was observed in the K18-hACE2 SARS-CoV2 infection mouse model.[11] Until now the SARS-CoV2 3CLpro remains to be validated as a promising therapeutic target; which is only possible with a 3CLpro inhibitor that lacks activity against other proteases such as cathepsin L. We applied a structure-based strategy with early monitoring of cathepsin L inhibition in parallel with SARS-CoV-2 3CLpro inhibition. As a result, we identified ALG-097111, a potent and selective SARS-CoV-2 3CLpro inhibitor, which has demonstrated proof of concept by inhibiting SARS-CoV-2 infection in an *in vivo* SARS-CoV-2 hamster model.

## Materials and Methods

### SARS-CoV-2 3CLpro and human cathepsin L biochemical assays

The SARS-CoV-2 3CLpro and human cathepsin L assays were performed as previously described [12].

### Human β-coronavirus OC43 assay

The human beta-coronavirus OC43 assay in HeLa cells was performed as previously described. [13]

### Human α-Coronavirus 229E assay

The human alpha-coronavirus 229E was purchased from Virapur (San Diego, CA) and propagated using MRC-5 human lung fibroblast cells (ATCC). Huh7 cells (JCRB cell Bank) were cultured using DMEM media, supplemented with 10 % fetal bovine serum (FBS), 1% (v/v) penicillin/streptomycin (P/S), 1% (v/v) HEPES and 1% (v/v) cellgro glutagro™ supplement (all Corning, Manassas, VA) at 37°C. 1.5 x 10^4^ Huh7 cells per well were cultured for 24 hours prior to infection. Then, serially diluted compounds in assay media (DMEM, 4% FBS, 1% P/S, 1% cellgro glutagro™ supplement, 1% HEPES) were added to the cells and incubated for 4 hours at 37°C. The 229E virus stock was diluted to a concentration known to produce optimal cytopathic effect and was added to cells in 96-well plates that were then incubated for 4 days at 33°C. Cellular cytotoxicity plates without the addition of 229E virus were set up in parallel. At the end of the incubation period, 100 μl of cell culture supernatant was replaced with 100 μl CellTiter-Glo® (Promega, Madison, WI) and incubated for at least 10 min at room temperature prior to measuring luminescence on a Perkin Elmer (Waltham, MA) Envision plate reader.

### SARS-CoV-2 nanoluciferase assay in human ACE-2 expressing A549 cells

The SARS-CoV-2 nanoluciferase assay using A549 cells expressing the human ACE-2 receptor was performed in the laboratory of Pei-Yong Shi at the University of Texas, Medical Branch [14] on behalf of Aligos Therapeutics, Inc. Cytotoxicity was assessed on A549 cells (Sigma) that were maintained in F12 Media, supplemented with 10% fetal bovine serum (FBS), 1% (v/v) penicillin/streptomycin (P/S), 1% (v/v) HEPES and 1% (v/v) cellgro glutagro™ supplement at 37°C with 5% CO2. 1.2 x 10^4^ A549 cells per well were plated in 96-well plates and cultured for up to 24hr. Compounds were diluted in F12 Media containing low FBS (2%) and added to cells and incubated for 48h. Reduction of cell viability was measured as described above.

#### Compounds

Ritonavir was purchased from Fluorochem (CAS 155213-675, FCC3125867). EIDD-2801 was purchased from Excenen Pharmatech Co., Ltd. GS-441524 (parent nucleoside of remdesivir) was obtained from Carbosynth (cat no AG167808). Compounds in table 1 were prepared via methodologies described in the literature. [15], [16], [17], [18], [19], [6]

**Table 1.** Overview of a set of reference compounds and their inhibitory potency on Cathepsin L and SARS-CoV-2 3CLpro in a biochemical assay.

|  | Cpd 11a<br>[15] | 6j<br>[16] | 6e<br>[16] | Cpd 13b<br>[17] | Cpd 11r<br>[18] | A9<br>[19] | PF-231<br>[6] |
| --- | --- | --- | --- | --- | --- | --- | --- |
| Cathepsin L<br>IC<sub>50</sub> (nM) | 0.21 (n=2) | <0.5 (n=2) | <0.5 (n=2) | 291 (n=2) | <0.5 (n=2) | <0.5 (n=2) | 146 (n=13) |
| SARS-CoV-2<br>3CLpro IC<sub>50</sub> (nM) | 8 (n=28) | 7 (n=2) | 10 (n=3) | 472 (n=2) | 1154 (n=2) | 4891 (n=2) | 5 (n=5) |

### SARS-CoV-2 infection model in Primary Human Lung Epithelial cells

Human small airway epithelia cell cultures were derived from a healthy donor and differentiated in an air-liquid culture system and as such obtained from Epithelix (cat no EP21SA). Cultures were maintained as described by the manufacturer before use. At the start of the experiment the apical side of the cultures were washed once with PBS and the inserts were transferred to wells containing the appropriate compound concentration in basal medium (SmallAir culture medium Epithelix). After 1.5 h incubation at (35°C, 5%CO2) the inserts were infected by adding SARS-CoV-2-GHB-03021/2020 at 2×10^4 TCID50 in 100 μL of PBS at the apical side. After 2 h incubation the virus inoculum was removed and the inserts were further incubated (35°C, 5%CO2). The basal medium (with or without compound) was replaced every two days. Virus production at the apical side was determined on day 1, 2, 4 and 6 by washing this side with 100 μL PBS and quantification of the amount of vRNA by first mixing 5 μL wash fluid with 50 μL Cells-to-cDNA II Cell Lysis Buffer (Thermofisher) and heating at 75°C or 15 minutes. The samples were subsequently diluted by adding 150 μL of water and 4 μL was used for RT-qPCR performed on a LightCycler96 platform (Roche) using the iTaq Universal Probes One-Step RT-qPCR kit (BioRad) with N1 primers and probes targeting the nucleocapsid (IDTDNA, cat no 10006770). Standards for the RT-qPCR were prepared by a 10-fold dilution of SARS-CoV-2-GHB-03021/2020 virus stock with known infectivity and using this in the same extraction/amplification protocol. The amount of vRNA in the samples could be calculated to TCID50 equivalents.

### SARS-CoV-2 infection model in hamsters

The hamster infection model of SARS-CoV-2, the SARS-CoV-2 isolate, SARS-CoV-2 RT-qPCR and End-point virus titrations has been described previously[20] [21].

### Treatment regimen

Female hamsters, 6-8 weeks old were anesthetized with ketamine/xylazine/atropine and inoculated intranasally with 50 μL containing 2×10^6^ TCID50 SARS-CoV-2-GHB-03021/2020 (day 0). Beginning 2h before infection, animals were treated twice daily with either vehicle used for ALG-097111 (60% PEG400 in water, subcutaneous[SC]) + ritonavir (50 mg/kg/dose, BID oral gavage) [negative control group] or ALG-097111 (200 mg/kg/dose, BID, SC) + ritonavir (50 mg/kg/dose, BID, oral gavage) or EIDD-2801 (200 mg/kg/dose, oral gavage as a positive control). Hamsters were monitored for appearance, behavior and weight. At day 2 post infection (pi) (day 3), 6 hours following the 5^th^ dose, hamsters were euthanized by IP injection of 500 μL Dolethal (200 mg/mL sodium pentobarbital, Vétoquinol SA). Lungs were collected and viral RNA and infectious virus were quantified by RT-qPCR and end-point virus titration, respectively.

### Pharmacokinetics (PK) Analysis of ALG-097111 in Plasma and Lung

#### Animal and treatments for PK evaluations

Male Sprague Dawley rats were group-housed during acclimation and in-life portions of the study. The animal room environment was controlled and monitored daily with target temperature 20 to 26°C, relative humidity 30 to 70%, 12 hours artificial light and 12 hours dark. All animals had access to certified rodent diet and reverse-osmosis water ad libitum. Animals were deprived of food overnight prior to dosing (fasted within 14 hours of dosing) and fed immediately after the 4-hour timepoint collection.

For the intravenous administration at 2 mg/kg, ALG-097111 was dissolved in 80% PEG400 in water. For the single dose administration via oral gavage at 10 mg/kg, ALG-097111 was dissolved in 40% PEG400 in water and dosed at 5 mL/kg. The formulation for subcutaneous administration at 50 mg/kg was 40% PEG400 in water. The blood collection in K2EDTA containing tubes was conducted at multiple timepoints from 0.083 to 24-hour postdose from both PO and SC dosed animals. The samples were kept cold on wet ice until centrifugation to harvest plasma.

Female golden Syrian hamsters were group-housed during acclimatization and in-life portions of the study. The animal room environment was controlled for temperature, humidity, and light/dark cycle as described for rats. All animals had access to certified rodent diet and reverse-osmosis water ad libitum. The animals were dosed 10h apart subcutaneously with 200 mg/kg/dose BID of ALG-097111 in 60% PEG400 in water. An oral gavage dose of ritonavir at 50 mg/kg in 95% PEG400 and 5% copovidone was administered just prior to each subcutaneous dose of ALG-097111. Blood samples at multiple timepoints from 0.25-to 24-hours postdose were collected into tubes containing K2EDTA and kept on ice until processing to harvest plasma. Lung samples were collected at 24 h after the first dose, flash frozen on dry ice and stored at ≤70 ºC until processing and analysis, then homogenized with methanol-water (7:3) on wet ice. The homogenate was centrifuged, and the supernatant was used for bioanalysis.

All in vivo studies were approved by the IACUC at their respective institutions.

#### Bioanalysis of plasma and lung extract

The samples were analyzed by LC-MS/MS method using ACQUITY UPLC system (Waters Corporation, Milford, MA) and Sciex Triple Quad™ 6500+ LC-MS/MS System. The plasma samples processed by protein precipitation method were injected on ACQUITY UPLC BEH C18 1.7 μm 2.1 × 50 mm column. A gradient of 0.1% formic acid in water and acetonitrile at 0.6 mL/min was used. The mass spectrum was operated in positive ionization mode.

#### In vitro stability in microsomes and hepatocytes

Stability studies with ALG-097111 at 1 μM were performed in hamster, dog and human liver microsomes, and in dog and human hepatocytes. Details can be found in the supplementary information.

### Secondary *in vitro* pharmacology

ALG-097111 was profiled up to a concentration of 10 μM in the safety 44 CEREP panel, a panel of receptor, enzyme and uptake assays (44 total) and on a panel of 58 kinases. ALG-097111 was tested on a set of 9 proteases (Calpain 1, Caspase 2, Cathepsin B, Cathepsin D, Cathepsin L, Chymotrypsin, Elastase, Thrombin a, and Trypsin).

## Results and discussion

The entry of SARS-CoV-2 into the target cells is a process that can be mediated by multiple proteases including cysteine cathepsins B and L or the transmembrane protease serine 2 (TMPRSS2). [22,23] The cathepsin L inhibitor K117777, which lacks an inhibitory effect on the 3CLpro results in potent inhibition of SARS-CoV-2 in VeroE6, A549-ACE2 and HeLa-ACE2.[24]

Although lung tissue cells express both cathepsins and TMPRSS2, many cell-lines commonly used for screening, such as A549, lack TMPRSS2. [9] Strikingly, the potent antiviral effect of K117777 was abolished when TMPRSS2 was expressed in A549-ACE2. Off target activity of 3CLpro inhibitors could lead to an inaccurate assessment of the potential activity of a lead candidate in vivo. Indeed, the antiviral potency in cell-based assays will be overestimated when tested on cell lines not reproducing all entry pathways. Moreover, these non-selective inhibitors will have to compete in vivo for viral and host enzymes, likely leading to a further reduction of the antiviral potency. In addition, cysteine cathepsins are key regulators of both the innate and adaptive immunity to pathogens [25] [26]. In particular, Cathepsin L primes T-cells by processing the antigens in the lysosomal compartment of Antigen Presenting Cells (APC’s), by presenting the proteolytic degradation peptides on the MHC class II, [27] processing the MHC class II [28] and different Toll-like receptors (TLR3, −7 and −9) [29], [30] and by controlling the secretion of different cytokines [25] [26]. Therefore, 3CLpro inhibitors that lack selectivity toward cathepsin L, may potentially have counter-effective side effects by dampening the immune response against SARS-CoV-2 [31]. This may be expected to aggravate the already impaired T-cell immunity of COVID-19 patients who often suffer from lymphocytopenia, reduced expression of MHCII and pro-inflammatory cytokines.

The selectivity of a set of reported SARS-CoV-2 3Clpro inhibitors versus cathepsin L, was assessed in a biochemical assay (Table 1). The examples depicted indicate that mimicking the substrates glutamine in P1 of the inhibitor, is not sufficient to obtain a selective inhibitor against the 3CL protease and that Cathepsin L tolerates this substituent, resulting in potent inhibition.

### In Vitro Profiling of ALG-097111

Structure-based optimization, combined with cathepsin L and SARS-CoV-2 3Clpro biochemical activities, resulted in the discovery of the selective 3Clpro inhibitor ALG-097111. In biochemical assays, ALG-097111 potently inhibited SARS-CoV-2 3CLpro (IC_50_ = 7 nM, n=5) without affecting the activity of human cathepsin L (IC_50_ > 10 μM, n=3) (Figure S1). The selectivity of ALG-097111 was confirmed by *in vitro* testing of a panel of 44 receptors, 58 kinases (no inhibition was noted up to 10 μM) and 9 human proteases (<50% inhibition at 10 μM).

In comparison, GC376 and PF-00835231 were significantly less selective against human cathepsin L. ALG-097111 inhibited replication of SARS-CoV-2 in A549 cells expressing the human ACE-2 receptor with an EC50 of 200 ± 18.4 nM (n=2). No cytotoxicity was observed in A549 cells at concentrations up to 100 μM, resulting in an in vitro selectivity of over 500. The activity of ALG-097111 extended to other human coronaviruses such as the alphacoronavirus 229E and the beta-coronavirus OC43, demonstrating broad-spectrum anti-coronaviral activity. (Table 2).

**Table 2:** Antiviral activity and cytotoxicity of ALG-097111, and remdesivir in various cell-based antiviral assays.

| | SARS-CoV-2Nluc<br>(A549)* | | β-CoV OC43<br>(HeLa) | | α-CoV 229E<br>(Huh-7) | |
| --- | --- | --- | --- | --- | --- | --- |
|  | EC<sub>50</sub><br>[nM] | CC<sub>50</sub><br>[nM] | EC<sub>50</sub><br>[nM] | CC<sub>50</sub><br>[nM] | EC<sub>50</sub><br>[nM] | CC<sub>50</sub><br>[nM] |
| ALG-097111 | 200 ± 18.4<br>(n=2) | > 100 000 | 123.3 ± 21.1<br>(n=4) | >100 000 | 366.5 ± 199.1<br>(n=2) | >25 000 |
| Remdesivir | 31.6 ± 11.9<br>(n=2) | > 50 000 | 102.4 ± 50.53<br>(n=46) | > 5000 | 14.47 ± 3.48<br>(n=18) | > 1000 |

\* the SARS-CoV-2-Nluc antiviral assay was performed using A549-hACE2 cells while the cytotoxicity was performed in A549 cells.

The anti-SARS-CoV-2 activity of ALG-097111 was confirmed in human small airway epithelial cell cultures derived from a healthy donor and differentiated in an air-liquid culture system (Figure S2). When ALG-097111 was added at a concentration of 1μM to the basolateral side of the cultures, the compound reduced viral RNA yield in the washes at the apical site of the culture by >3 log_10_.

While ALG-097111 is stable in human and dog liver microsomes (t1/2 >60 min) and hepatocytes (t1/2 >360 min), it has a lower stability in hamster liver microsomes (t1/2= 15 min). In the presence of ritonavir (μM), the in vitro half-life in hamster liver microsomes increased (t1/2= > 60 min)

### In vivo Pharmacokinetic evaluation of ALG-097111

ALG-097111 displayed low clearance (11.2 mL/min/kg), moderate volume of distribution (0.79 L/kg) and half-life of 2.0 h following a single intravenous dose at 2 mg/kg in rats. Plasma PK profile of ALG-097111 in rats was obtained following a single dose at 10 mg/kg administered via oral gavage and at 50 mg/kg via subcutaneous injection. Although, the bioavailability following the oral dose was low (3.2%). The systemic exposure following the subcutaneous administration was significantly higher with bioavailability of 44%. The comparative plasma profile is depicted in Figure 1A.

**Figure 1:**
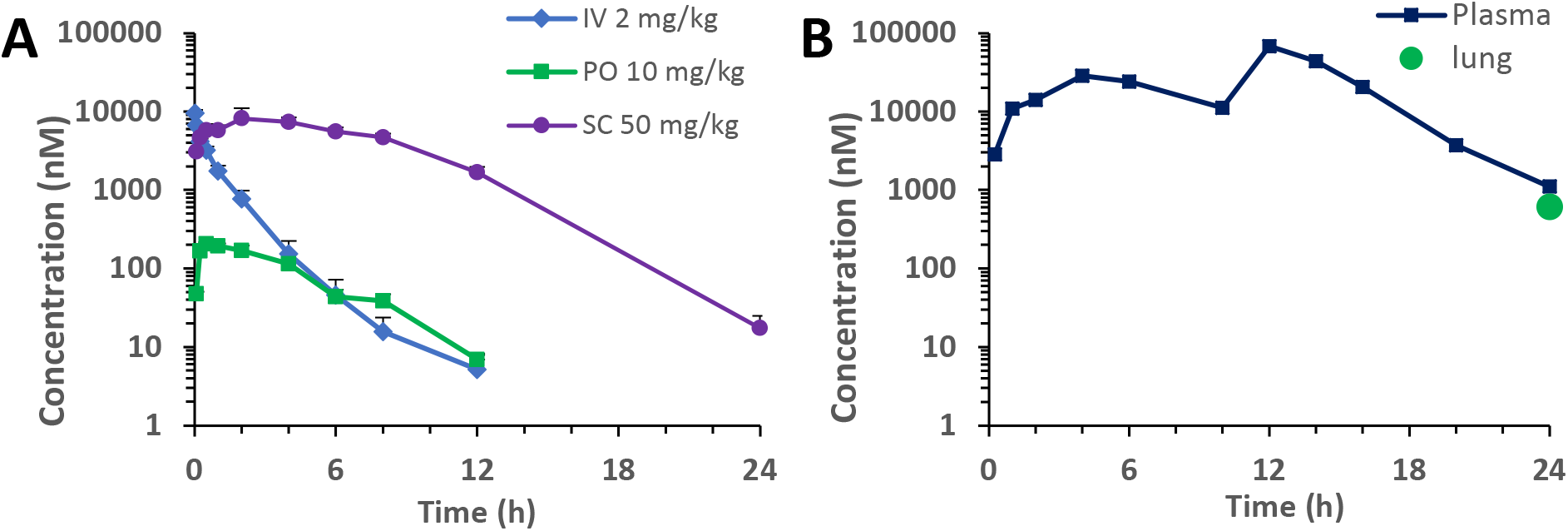
A) Plasma PK of ALG-097111 in male Sprague Dawley rats following a single IV, PO or SC dose. B) Plasma PK profile and lung concentration at 24 h of ALG-097111 in female golden Syrian hamsters following BID administration of ritonavir given orally at 50 mg/kg/BID prior to ALG-097111 given subcutaneously at 200 mg/kg/dose.

When ALG-097111 was co-administered in female hamsters at 200 mg/kg/dose BID with ritonavir given PO at 50 mg/kg/dose BID, the plasma and lung Ctrough of 1105 nM and 611 nM, were 5.5 and 3.0-fold above the *in vitro* EC50 on SARS-CoV-2 in A549, respectively. The hamster plasma PK profile and lung concentration at 24 h are shown in Figure 1B.

### Evaluation of In Vivo Efficacy of ALG-097111 in SARS-CoV-2 infected Syrian Hamsters

The high *in vitro* efficacy of the compound for inhibiting SARS-CoV-2 virus combined with its *in vitro* metabolic stability in nonrodents and human matrices and PK profile in rat led to selection of the compound for *in vivo* evaluation of its efficacy. The in vivo anti-SARS-CoV-2 efficacy of ALG-097111 was evaluated in female SG hamsters dosed twice daily with 200 mg/kg of ALG-097111 in combination with ritonavir (50 mg/kg/dose). Other hamsters were treated orally BID with the vehicle+ritonavir (i.e. the control group) or molnupiravir (EIDD-2801) (200 mg/kg as a positive control group). All animals received 5 administrations in total, starting 2h before infection. At day 2 pi, the animals were euthanized, and lungs were collected for quantification of viral loads. Treatment of infected hamsters with ALG-097111 resulted in 3.5 log_10_ reduction in the viral RNA copies per mg of lung tissue (P=0.008) as compared to the vehicle/ritonavir-treated animals (Figure 2A). In hamsters that had been treated with 200 mg/kg BID of molnupiravir (EIDD-2801) a 2.0 log_10_ reduction in the viral RNA copies/mg of lung tissue was noted (P=0.095, non-significant) (Figure 2A). Both compounds significantly reduced the infectious virus titers in the lungs by 3.7 (P=0.008) and 4.1 (P=0.008) log_10_ TCID50 per mg for ALG-097111 and EIDD-2801, respectively (Figure 2B).

**Figure 2.**
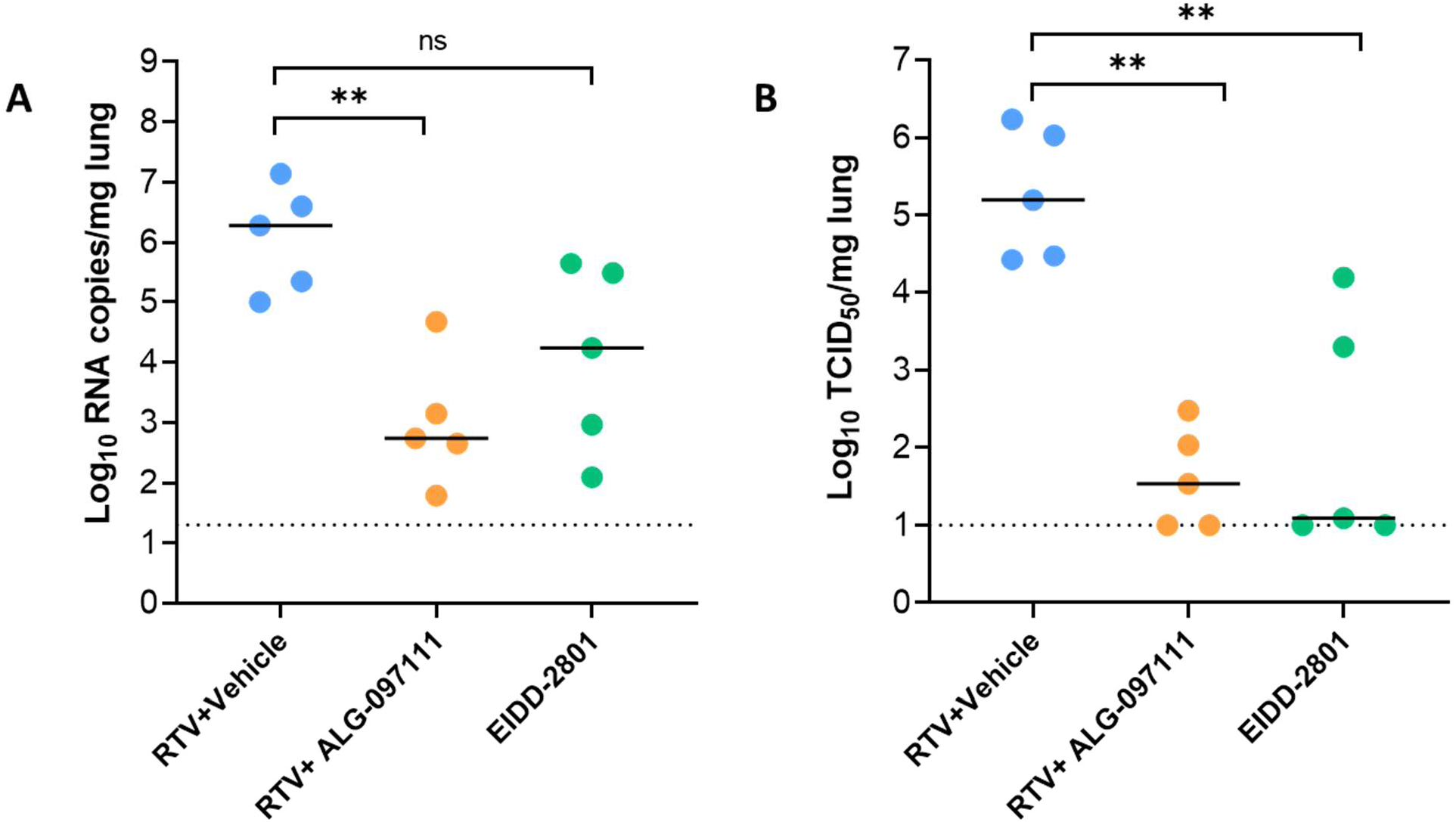
*In vivo* efficacy of ALG-097111 against SARS-CoV-2 in SG hamsters. (A) Viral RNA levels in the lungs of ritonavir (PO)+vehicle (SC) (50 +0 mg/kg/dose, BID), ritonavir (PO)+ALG-097111 SC (50 +200 mg/kg/dose, BID, SC) and EIDD-2801-PO (200 mg/kg/dose, BID) SARS-CoV-2-infected hamsters at day 2 postinfection (pi) are expressed as log_10_ SARS-CoV-2 RNA copies per mg lung tissue. Individual data and median values are presented. (B) Infectious viral loads in the lungs of ritonavir (PO)+vehicle SC (50+ 0 mg/kg/dose, BID), ritonavir (PO) +ALG-097111 SC (50+200 mg/kg/dose, BID) and EIDD-280 (PO) (200 mg/kg/dose, BID) SARS-CoV-2-infected hamsters at day 2 post-infection (pi) are expressed as log_10_ TCID50 per mg lung tissue Individual data and median values are presented. Data were analyzed with the Mann-Whitney U test. **P < 0.01, ns=non-significant. RTV=ritonavir.

In conclusion, ALG-097111, a selective SARS-CoV-2 3CLpro inhibitor, is a potent inhibitor of the *in vitro* replication of SARS-CoV-2 and efficiently inhibits viral replication in the lungs of infected hamsters. To the best of our knowledge, this is the first time that a selective SARS-CoV-2 3CLpro inhibitor (i.e. without a significant inhibitory effect on cathepsin L activity), has been shown to result in such a pronounced inhibitory effect on SARS-CoV-2 replication in an animal infection model. Future studies will focus on combinations with other modalities against SARS-CoV-2, as well as investigation of compound dosing in a therapeutic setting.

## Acknowledgments

We thank Carolien De Keyzer, Lindsey Bervoets, Thibault Francken, Birgit Voeten, Niels Cremers, Tina Van Buyten, Joost Schepers, Winston Chiu and Kim Donckers for excellent technical assistance. We are grateful to Piet Maes for kindly providing the SARS-CoV-2 strain used in this study. A special thanks to Dr Hongjie Xia in the laboratory of Prof Dr Pei-Yong Shi at the University of Texas Medical Branch for kindly performing the SARS-CoV-2 Nanoluc antiviral assay.

## References

[1] A.A. Agbowuro, et al.,, Proteases and protease inhibitors in infectious diseases, Medicinal Research Reviews 38 (2018) 1295–1331. http://doi.org/10.1002/med.21475.

[2] W. Zhu, et al., RNA-Dependent RNA Polymerase as a Target for COVID-19 Drug Discovery, SLAS DISCOVERY: Advancing the Science of Drug Discovery 25 (2020) 1141–1151. https://doi.org/10.1177/2472555220942123.

[3] Y. Wang, et al., Remdesivir in adults with severe COVID-19: a randomised, double-blind, placebo-controlled, multicentre trial, The Lancet 395 (2020) 1569–1578. https://doi.org/10.1016/S0140-6736(20)31022-9.

[4] Efficacy and Safety of Molnupiravir (MK-4482) in Hospitalized Adult Participants With COVID-19 (MK-4482-001), https://ClinicalTrials.gov/show/NCT04575584.

[5] Efficacy and Safety of Molnupiravir (MK-4482) in Non-Hospitalized Adult Participants With COVID-19 (MK-4482-002), https://ClinicalTrials.gov/show/NCT04575597.

[6] R.L. Hoffman, et al., Discovery of Ketone-Based Covalent Inhibitors of Coronavirus 3CL Proteases for the Potential Therapeutic Treatment of COVID-19, Journal of Medicinal Chemistry 63 (2020) 12725–12747. https://doi.org/10.1021/acs.jmedchem.0c01063.

[7] L. Hoffman Robert, et al., Anticoronviral Compounds And Compositions, Their Pharmaceutical Uses And Materials For Their Synthesis, Pfizer, WO2005113580.

[8] C.-C. Chen, et al., Overview of antiviral drug candidates targeting coronaviral 3C-like main proteases, The FEBS Journal n/a (2021). https://doi.org/10.1111/febs.15696.

[9] K. Steuten, et al., Challenges for targeting SARS-CoV-2 proteases as a therapeutic strategy for COVID-19, bioRxiv (2020) 2020.2011.2021.392753. https://doi.org/10.1101/2020.11.21.392753.

[10] Y. Shi, et al., The Preclinical Inhibitor GS441524 in Combination with GC376 Efficaciously Inhibited the Proliferation of SARS-CoV-2 in the Mouse Respiratory Tract, bioRxiv (2020) 2020.2011.2012.380931. https://doi.org/10.1101/2020.11.12.380931.

[11] C. Joaquín Cáceres, et al., Efficacy of GC-376 against SARS-CoV-2 virus infection in the K18 hACE2 transgenic mouse model, bioRxiv (2021) 2021.2001.2027.428428. https://doi.org/10.1101/2021.01.27.428428.

[12] C. Liu, et al. Dual inhibition of SARS-CoV-2 and human rhinovirus with protease inhibitors in clinical development, Antiviral Res 187 (2021) 105020. 10.1016/j.antiviral.2021.105020.

[13] Z.A. Gurard-Levin, et al., Evaluation of SARS-CoV-2 3C-like protease inhibitors using self-assembled monolayer desorption ionization mass spectrometry, Antiviral Research 182 (2020) 104924. https://doi.org/10.1016/j.antiviral.2020.104924.

[14] X. Xie, et al., A nanoluciferase SARS-CoV-2 for rapid neutralization testing and screening of anti-infective drugs for COVID-19, Nature Communications 11 (2020) 5214. https://doi.org/10.1038/s41467-020-19055-7.

[15] W. Dai, et al. Structure-based design of antiviral drug candidates targeting the SARS-CoV-2 main protease, Science 368 (2020) 1331. https://doi.org/10.1126/science.abb4489.

[16] A.D. Rathnayake, et al., 3C-like protease inhibitors block coronavirus replication in vitro and improve survival in MERS-CoV-infected mice, Science Translational Medicine 12 (2020) eabc5332. https://doi.org/10.1126/scitranslmed.abc5332.

[17] L. Zhang, et al., Crystal structure of SARS-CoV-2 main protease provides a basis for design of improved α-ketoamide inhibitors, Science 368 (2020) 409. https://doi.org/10.1126/science.abb3405.

[18] L. Zhang, et al., α-Ketoamides as Broad-Spectrum Inhibitors of Coronavirus and Enterovirus Replication: Structure-Based Design, Synthesis, and Activity Assessment, Journal of Medicinal Chemistry 63 (2020) 4562–4578. 10.1021/acs.jmedchem.9b01828.

[19] H. Liu, et al., Ketoamide Compound And Preparation Method, Pharmaceutical Composition, And Use Thereof, Shanghai inst materia medica; univ fudan, WO2020030143.

[20] R. Boudewijns, et al., STAT2 signaling restricts viral dissemination but drives severe pneumonia in SARS-CoV-2 infected hamsters, Nature Communications 11 (2020) 5838. https://doi.org/10.1038/s41467-020-19684-y.

[21] S.J.F. Kaptein, et al., Favipiravir at high doses has potent antiviral activity in SARS-CoV-2-infected hamsters, whereas hydroxychloroquine lacks activity, Proceedings of the National Academy of Sciences 117 (2020) 26955. 10.1073/pnas.2014441117.

[22] J. Shang, et al., Cell entry mechanisms of SARS-CoV-2, Proceedings of the National Academy of Sciences 117 (2020) 11727. https://doi.org/10.1073/pnas.2003138117.

[23] M. Hoffmann, et al., SARS-CoV-2 Cell Entry Depends on ACE2 and TMPRSS2 and Is Blocked by a Clinically Proven Protease Inhibitor, Cell 181 (2020) 271-280.e278. https://doi.org/10.1016/j.cell.2020.02.052.

[24] D.M. Mellott, et al., A cysteine protease inhibitor blocks SARS-CoV-2 infection of human and monkey cells, bioRxiv (2020) 2020.2010.2023.347534. https://doi.org/10.1101/2020.10.23.347534.

[25] P.I. Bird, et al., Endolysosomal proteases and their inhibitors in immunity, Nature Reviews Immunology 9 (2009) 871-882. 10.1038/nri2671.

[26] T. Jakoš, et al., Cysteine Cathepsins in Tumor-Associated Immune Cells, Frontiers in Immunology 10 (2019). 10.3389/fimmu.2019.02037.

[27] J.D. Colbert, et al., Diverse regulatory roles for lysosomal proteases in the immune response, European Journal of Immunology 39 (2009) 2955–2965. https://doi.org/10.1002/eji.200939650.

[28] T. Yadati, et al., The Ins and Outs of Cathepsins: Physiological Function and Role in Disease Management, Cells 9 (2020). https://doi.org/10.3390/cells9071679.

[29] S.E. Ewald, et al., Nucleic acid recognition by Toll-like receptors is coupled to stepwise processing by cathepsins and asparagine endopeptidase, Journal of Experimental Medicine 208 (2011) 643–651. https://doi.org/10.1084/jem.20100682.

[30] A. Garcia-Cattaneo, et al., Cleavage of Toll-like receptor 3 by cathepsins B and H is essential for signaling, Proceedings of the National Academy of Sciences 109 (2012) 9053. https://doi.org/10.1073/pnas.1115091109.

[31] M.V. Baranov, et al., The PIKfyve Inhibitor Apilimod: A Double-Edged Sword against COVID-19, Cells 10 (2021). https://doi.org/10.3390/cells10010030.

