## Supplemental information for "ALG-097111, a potent and selective SARS-CoV-2 3-chymotrypsin-like cysteine protease inhibitor exhibits *in vivo* efficacy in a Syrian Hamster model"

### Content

Page 2: Figure S1

Page 3: Figure S2

Page 4: Protocol of *in vitro* stability in microsomes and hepatocytes

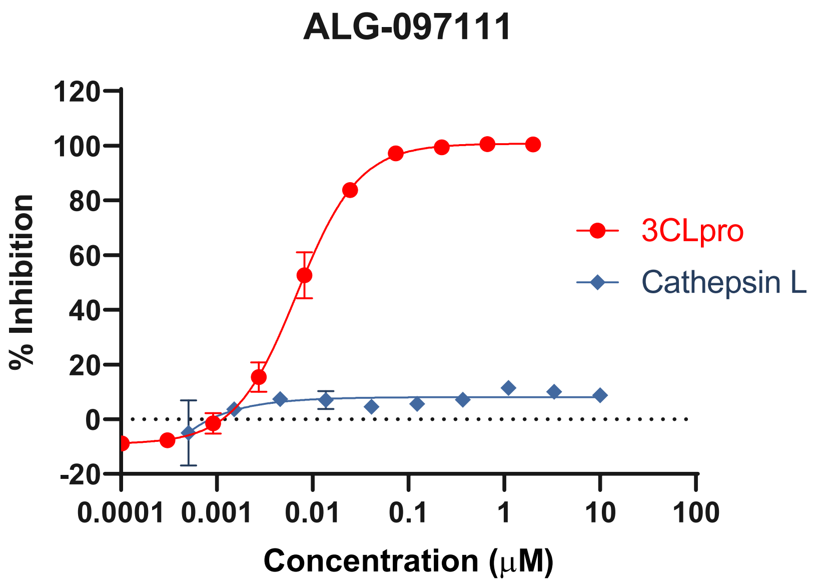


Figure S1. Biochemical activity of ALG-097111 on Cathepsin L (Blue) and 3CLpro (red).

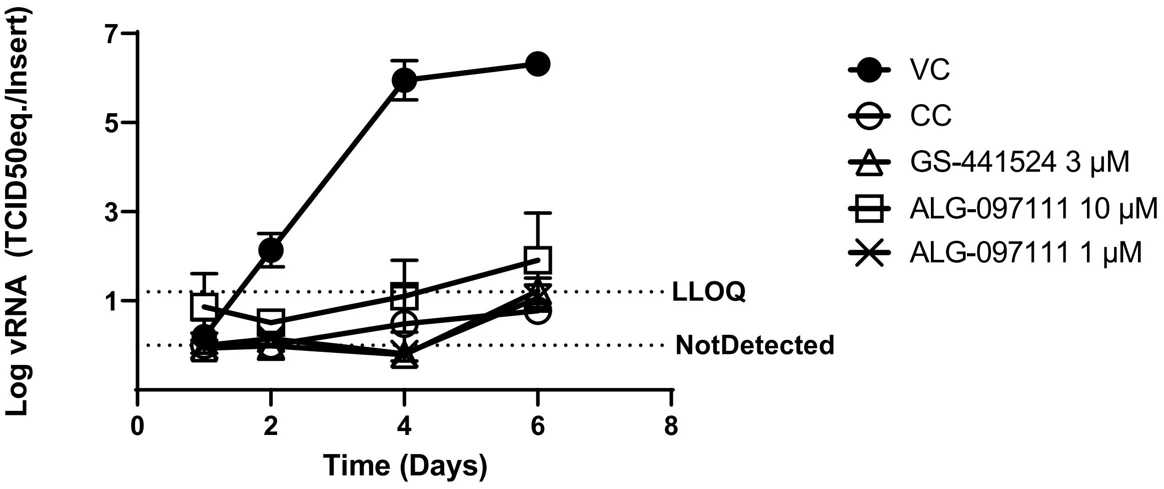


Figure S2: Anti-SARS-CoV-2 activity in primary human lung epithelial cell cultures. Human small airway epithelia cells, cultivated and differentiated in an air-liquid system, were treated from the basal site with the indicated concentrations of compound starting 1.5 h before infection with SARS-CoV-2-GHB-03021/2020 (2x10^4 $TCID_{50}$/insert). Viral RNA production on the apical site was quantified at different time points by RT-qPCR. Each condition was tested in 3 independent cultures and averages and STDEV are shown. 'LLOQ' indicates the lower limit of quantification and is defined by the point in the linear part of the standard curve with the lowest vRNA concentration. 'Not Detected' relates to a CT-value of > 36.

In vitro stability in microsomes and hepatocytes

ALG-097111 was incubated at 1 μM at 37 °C with 0.5 mg/mL of hamster, dog, human liver microsomes for 0, 15, 30 and 60 minutes. The incubation in hamster liver microsomes was conducted in the presence of ritonavir (1 μM) also. Verapamil at 1 μM was included as a positive control to verify suitability of the assay system. It was incubated at 1 μM at 37 °C in cryopreserved hepatocytes from dog and human at 0.5 million cells/mL for 0, 60, 120 and 180 minutes. Midazolam at 1 μM was included as a positive control. The samples were analyzed by LC/MS/MS. The rates of clearance of test compound were calculated using linear regression plot of semi-log % remaining of the test compound versus time. The elimination rate constant of the linear regression plot was then used to determine half-life.
